# On the Analysis of Transcriptional Noise From RNA-sequencing Data

**DOI:** 10.1101/2021.04.06.438605

**Authors:** Kristoffer Vitting-Seerup

## Abstract

RNA-sequencing (RNA-seq) has revolutionized our understanding of molecular and cellular biology. A central cornerstone in the analysis of RNA-seq is the bioinformatic tools that quantify the data. To evaluate the efficacy of these tools, scientists rely heavily on simulation of RNA-seq. Recently Varabyou *et al*. took simulation of RNA-seq data to the next level by providing simulated data, that includes simulation of transcriptional noise. While this represents a significant step forward in our ability to perform realistic benchmarks of RNA-seq tools, the data provided by Varabyou *et al*. need refinement. In the following, I suggest a few improvements with a specific focus on splicing noise.

**Preface:** I wrote this paper intending to submit it as a Commentary on the Varabyou *et al*. 2020 Genome Research paper^1^, but apparently, Genome Research does not publish correspondence-type articles. That is why it is currently on BioRxiv. If you have suggestions about where this paper could potentially be published do not hesitate to contact me.

## Introduction

RNA-sequencing (RNA-seq) has revolutionized our understanding of molecular- and cell biology. It is one of the most widely used high throughput methods, with increasing amounts of data produced each year. It naturally follows that accurate quantification methods are paramount. Several bioinformatic methods have over the last decade revolutionized transcriptomics quantification by enabling accurate transcript-level quantification of RNA-seq^2–5^. An essential component in this development is the benchmarking of these tools. Such benchmarking relies heavily on the simulation of RNA-seq data^2–4^.

Recently Varabyou *et al*^1^. reported the first attempt at assessing the extent of transcriptional noise in RNA-seq data and include it in RNA-seq simulations. To identify transcriptional noise, Varabyou *et al*. re-use the data from the impressive CHESS project^6^. In the CHESS project, Pertea *et al*. use StringTie^4^ to do a guided transcript assembly on the GTEx dataset^7^, which contained 9795 human RNA-seq samples from 49 tissues. The result of running the StringTie pipeline on all these samples was, after some filtering, ~20.7 million distinct transcripts, of which ~11.8 million overlapped known genes^1^. Next, Pertea *et al*. applied a series of stringent filters that reduced the ~20.7 million transcripts to the ~300.000 that constitute the CHESS reference database^6^. In Varabyou *et al*., the authors define all the ~20.4 million transcripts that did not make it into the final CHESS reference as noise^1^. Depending on the extend and type of overlap with the CHESS database, these noise transcripts were further sub-classified as intergenic, intronic, and splice noise. Varabyou *et al*. then sampled a set of real and noise transcripts and used the Polyester tool^8^ to simulate the noise containing RNA-seq data use for the their benchmark.

While this research represents a significant step forward in our ability to benchmark RNA-seq quantification methods, the analysis and simulated data provided by Varabyou *et al*. suffer from a series of potential problems that I will describe in the following.

## Results

### The Overlap of Transcript Types

To select what transcripts are provided to Polyester, Varabyou *et al*. use an elaborate sampling scheme based on quantifying both reference and noise transcripts in the GTEx data. While this approach preserves inherent transcription relationships between transcripts, it appears that it has an unintended consequence: Noise is simulated from a large number of genes where expression of real transcript is not simulated. The overlap between genes with real and splicing noise is shown for a representative sample in Figure 1A. Across all 30 simulations provided by Varabyou *et al*., I find, on average, only 43.5% of intronic noise and 53.7% of splicing noise originate from genes where expression of real transcripts is also simulated (Figure 1B). Wrong transcript assembly, splicing noise (and to some extent also intronic noise) is expected to be a by-product of real transcription^10^. Therefore, the lack of real transcripts could be problematic and might affect the false positive numbers and false positive rates reported in the benchmark presented by Varabyou *et al*.

**Figure 1:**
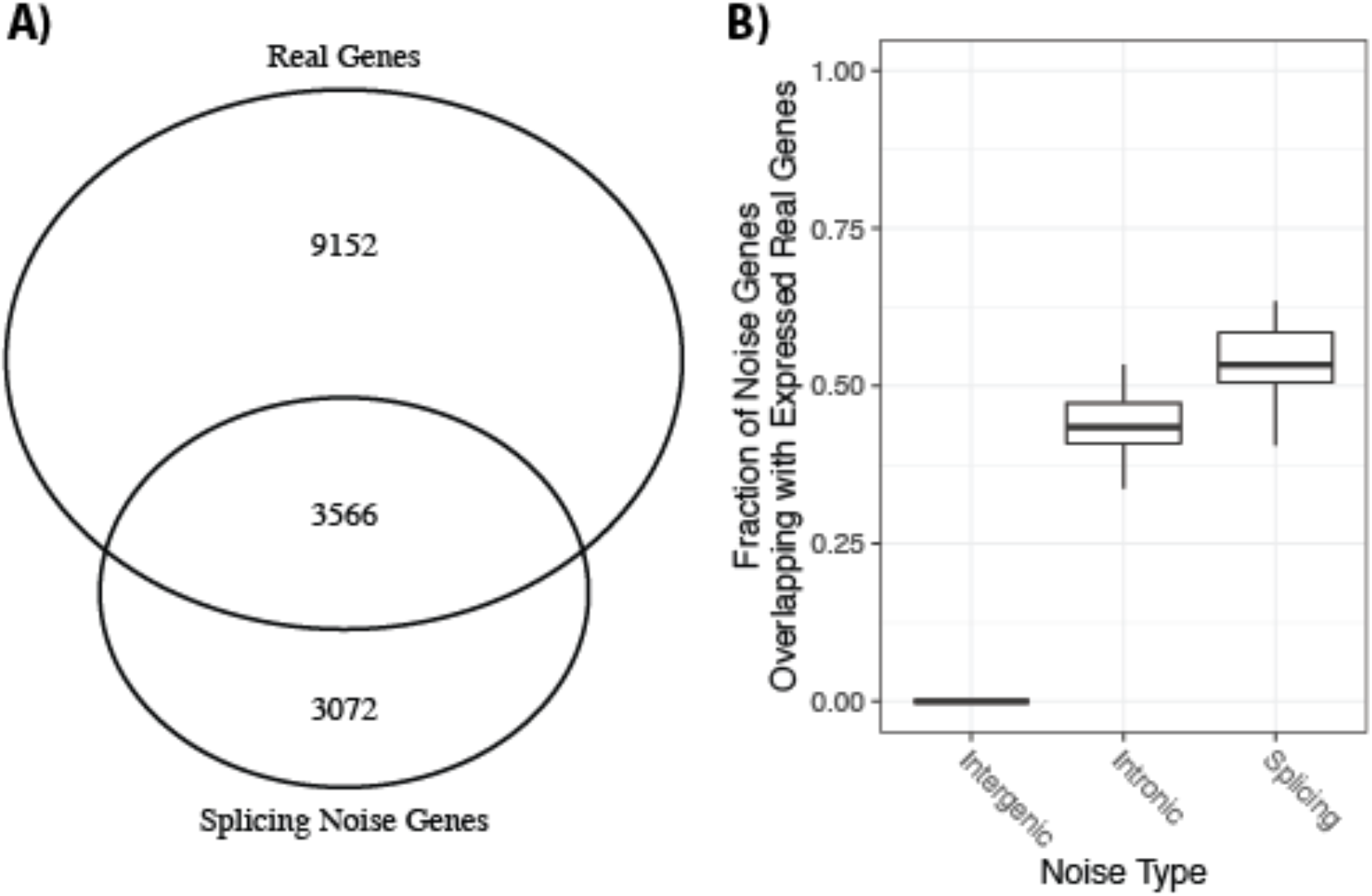
Overlap of genes containing real and noise transcripts. **A**) Venn diagram of genes with real and splicing noise from a representative sample (Tissue 1, sample 9). **B**) Summary of overlap in all 30 simulations. For each type of noise (x-axis), the fraction of genes with simulated noise overlaps with the simulation of real transcripts (y-axis).

### Defining Real and Noise Transcripts

Another potential problem arises due to the different objectives of the CHESS and Varabyou *et al*. studies. The aim of the CHESS project was to establish a new reference transcriptome. This task naturally necessitates the application of a long list of strict filters. In contrast, Varabyou *et al*. aim to estimate “realistic transcriptional noise”. For this aim, strict filtering could be problematic since many real transcripts would be defined as noise. In short, by directly reusing the CHESS data, Varabyou *et al*. might be too stringent in the definitions of which transcripts are real and which are noise thereby overestimating the noise.

To assess this problem from an orthogonal angle, I use IsoformSwitchAnalyzeR^11^ to characterize the transcript features of the Varabyou *et al*. simulations. From analyzing Open Reading Frames (ORF), the coding potential^12^, protein domains^13^, Premature Termination Codons (PTC), and the length of different transcript features, it is clear that intronic and intergenic noise transcripts are very different from real transcripts (Figure 2). On the other hand, a non-negligible fraction of the transcripts defined as splicing noise appears to be indistinguishable from real transcripts (Figure 2). The main difference between real and splicing transcripts seems to be that a smaller fraction (70.9% of real, 46.9% of splicing) have coding potential and that, in general, splice noise transcripts are somewhat shorter and have shorter ORFs. The shorter transcripts could be explained by the multiple CHESS filtering step where transcripts “contained in another transcript” are excluded^6^. However, this is speculation since it is unclear what “contained” means in the context of both transcript and exon boundaries (e.g., will a transcript with a downstream alternative donor splice site or an exon-skipping event be removed?). By classifying these seemingly real transcripts as noise, the noise estimates provided in Varabyou *et al*. could be inflated.

**Figure 2:**
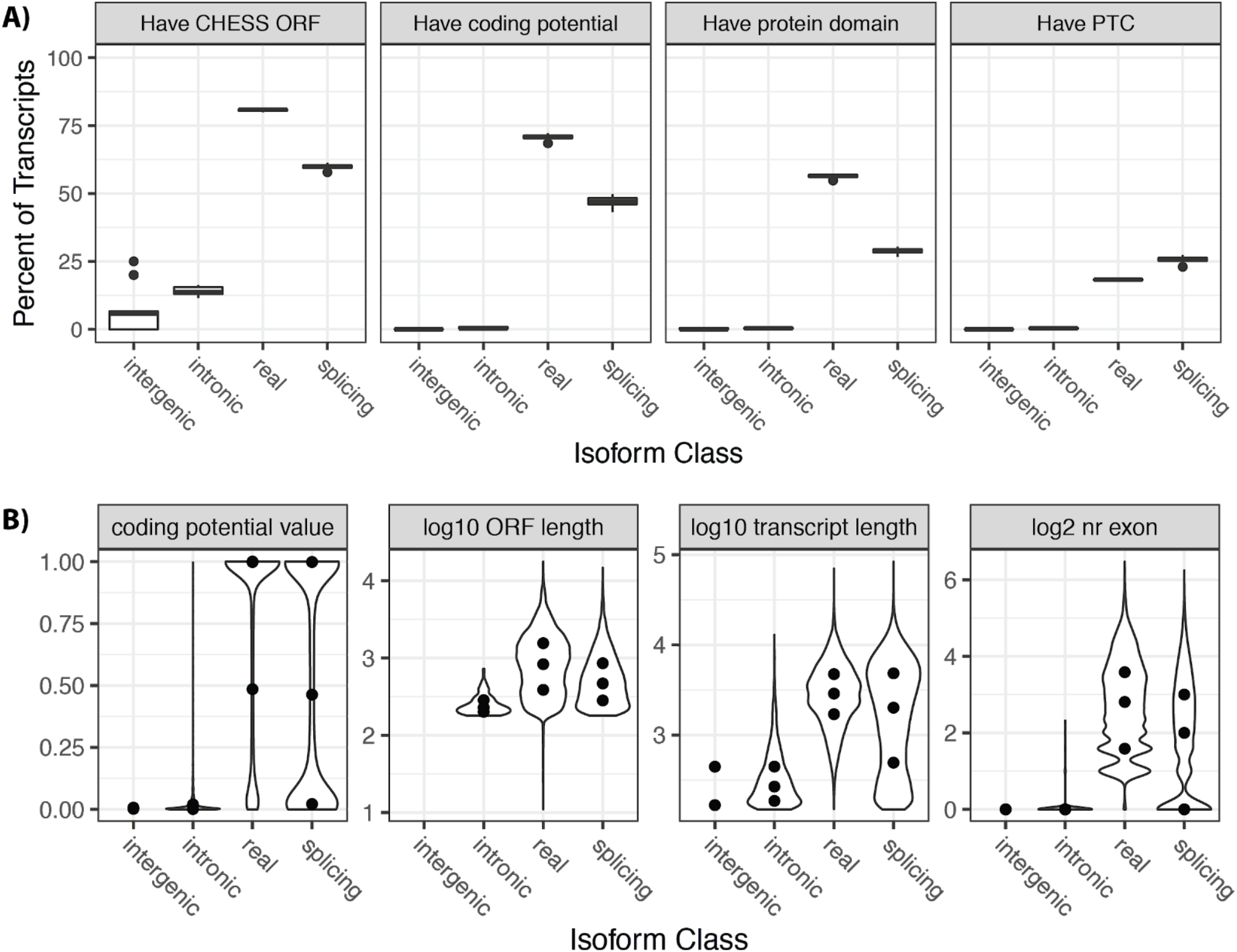
Characterization of transcripts included in the Varabyou *et al*. simulations. **A)** For each transcript type (x-axis), the percent of transcripts (y-axis) with specific annotations is shown. Subplots indicate different annotations. PTC: Premature Termination Codon (which typically results in the transcript being degraded by Nonsense Mediated Decay (NMD). Coding potential is calculated by CPAT. **B)** Overview of continuous variables for a representative sample. For each transcript type (x-axis), different continuous variables (sub-plots) are shown as violin plots where dots indicate 25th, 50th (median), and 75th percentile. Coding potential is calculated by CPAT.

### Choice of simulation type

Varabyou *et al*. choose to use Polyester^8^ to simulate single-end RNA-seq data. This seems strange as the quantification tools used are all 1) Designed to utilize paired-end data^2–4^ and 2) More accurate when used with paired-end data^14^. Varabyou *et al*. writes that it is due to problematic coverage of transcripts shorter than the fragment size. This argumentation seems peculiar given that most RNA-seq protocols specifically select for fragments transcripts in the 100-500 nucleotide range meaning this problem will only affect the less than 0.1% of the simulated “real” transcript that are shorter than 500 nucleotides. Notably the GTEx data used by the authors to both define and simulate noise transcripts was created using an RNA-seq protocol which includes a size-selection step.

### Aggregating Noise

Lastly a potential problem could arise from how the combined reference transcriptome was created. Varabyou *et al*. aggregate information from 9795 RNA-seq datasets to a single reference set containing both real and noise transcripts. All these 20.7 million transcripts (~486 transcripts per gene) are then used as a reference to quantify each individual GTEx sample. Noise levels are subsequently defined as the combined abundance of the transcripts defined as noise. Since transcript level quantification from short reads is challenging and affected by both the type of data^14^ (single vs paired-end) and the accuracy decreases as the number of transcripts increases^15^ the aggregation of noise from thousands of samples could inflate the noise estimates in individual samples simply because so many overlapping transcripts are quantified. Since the GTEx data is used to define the abundance levels for the RNA-seq simulation this problem will also have a spillover effect to the benchmark performed by Varabyou *et al*. Furthermore this problem probably increases when considering that the algorithm used to reconstruct the transcripts in the first place, like any other algorithm, is not perfect^4^, meaning assembly errors will also be accumulated and subsequently defined as noise.

## Discussion

While I believe including transcriptional noise in RNA-seq simulations is an important step forward, the approach used by Varabyou *et al*. leaves room for improvement. Here I have discussed potential improvement in the overlap of transcripts, the definition of “noise” vs. “real”, the choice of simulation type as well as the problem of aggregating. From this it is clear that simulating transcriptional noise is not straight-forward and there are still many unanswered questions, including determining the extend of (the different types of) transcriptional noise in humans.

## Data availability

The data collected via the IsoformSwitchAnalyzeR workflow for the representative sample (see methods) is summarized in the supplementary table and is available un-summarized via the four switchAnalyzeRlist R objects (saved as a single Rdata file) that can be found at doi.org/10.6084/m9.figshare.14307842. All data and scripts are available upon request.

## Methods

### Data

I downloaded the data Varabyou *et al*. used to simulate the RNA-seq data from https://doi.org/10.25739/v903-wd86 (The “simulated_experiments.tar.gz” file). The CHESS v2.2 reference GTF was downloaded from http://ccb.jhu.edu/chess/.

### Gene overlap

I imported each of the 120 simulated GTF (4 transcript type for each of the 30 simulations) files into R using rtracklayer::import() and extracted the sample name and transcript type (both from the filename) and gene ids (from the GTF file). Gene ids for different transcript types were compared within the same sample. The representable sample shown in figure 1A was chosen as the one where the overlap was closest to the average (t1_s9).

### Transcript analysis with IsoformSwitchAnalyzeR

I used IsoformSwitchAnalyzeR [Ref] 1.13.06 for all analyses modifying the entire workflow to not only analyze isoform switches by setting onlySwitchingGenes = FALSE in all functions.

From each of the 120 simulated GTFs, I created a switchAnalyzeRlist by importing the GTF into R using rtracklayer::import(), creating a dummy count matrix and design matrix which allows us to use the importRdata() function. In addition to the isoforms from the simulated GTF, I added all CHESS 2.2. reference transcripts from the same genes which were not already present in the switchAnalyzeRlists were created.

CDS were added as ORFs for transcripts originating from the CHESS reference using the addORFfromGTF() function. Next, I analyzed all transcripts not already annotated with an ORF for ORFs using the analyzeNovelIsoformORF() function with the analysisAllIsoformsWithoutORF = TRUE, orfMethod = ‘longest.AnnotatedWhenPossible’ and minORFlength = 180 arguments. This means that all transcripts without an annotated ORF were analyzed for potential ORFs. If the transcripts overlapped known translation start sites, these were used for the ORF analysis. An ORF had to be ORF to be at least 180 nucleotides (60 amino acids) to keep the same (stringent) criteria as the original CHESS analysis. To distinguish it from the more standard ORF definition of 100 nucleotides (33-34 amino acids), I refer to these longer ORFs as CHESS ORFs.

The biological sequences of the transcript sequences were extracted with the extractSequence() function. For both amino acids and nucleotide sequences, I created one non-redundant fasta file containing the sequences from all switchAnalyzeRlists. Coding potential was analyzed with CPAT v 1.2.3 [Ref], and protein domains were analyzed using Pfam [Ref].

The coding potential analysis was added to the switchAnalyzeRlists using analyzeCPAT() using the codingCutoff = 0.725 and removeNoncodinORFs = FALSE arguments. Protein domains were added to the switchAnalyzeRlist using analyzePFAM().

To analyze alternative splicing, the switchAnalyzeRlists were first reduced to multi-transcript genes using preFilter() and analyzing splicing with the analyzeAlternativeSplicing() function.

Complete data and summary statistics were extracted, calculated, and visualized using tidyverse. The representable sample shown in Figure 2B was chosen based on which sample had the fraction of transcripts with CHESS ORFs closes to the mean of all samples (t0_s5).

